# Unveiling the time course of visual stabilization through human electrophysiology

**DOI:** 10.1101/2022.12.08.519648

**Authors:** Yong Hoon Chung, Viola S. Stoermer

**Affiliations:** Department of Psychological and Brain Sciences, Dartmouth College

## Abstract

Positions of objects are coded relative to their surroundings, presumably providing visual stability even when we move our eyes around a visual scene. But when does this perceived stability of objects arise? Here we use a visual illusion, the frame-induced position shift, and measure electrophysiological activity elicited by an object whose perceived position is either shifted due to a surrounding frame or not, thus dissociating perceived and physical locations. We found that early visually-evoked responses were sensitive to physical location information earlier in time (∼70ms) than perceived location information (∼140ms). Furthermore, we show that location information can be reliably decoded across physical and perceived locations during the later time interval (140-180ms) but not during the earlier time interval (70-110ms). Together these results suggest that visual stability of objects emerges relatively late and is thus dependent on recurrent feedback from higher processing stages.

## Introduction

In everyday life, we constantly encounter dynamic visual inputs, especially due to frequent eye movements. However, despite the constant shifts on our retinas, the world appears mostly stable to us. One way to deal with such dynamicity in visual inputs is to code the information relative to the surrounding environment as the relational information between visual stimuli is often much more stable than the location information of an individual stimulus. For instance, when looking at a book on top of a table, the relational coding of the location of the book and the table will not change even when moving your eyes and shifting the retinal locations of the two objects. This allocentric processing has been shown to influence visual perception in a variety of ways, such as perceiving the direction and velocity of object motion (Post, Welch, & Whitney, 2008; Wertheim, 1981), the sense of heading direction (Morgan et al., 2015), and processing of object orientation (Asch & Witkin, 1948).

One recent study by Özkan et al. (2021) demonstrates a particularly strong example of how relational position coding can affect perception. This so-called Frame Induced Position Shift (FIPS) illusion elicits a paradoxical stabilization of a moving object: When a frame is moved left to right and two probe stimuli are flashed inside the frame at the exact same physical location shortly before and after the frame moves, participants report these probes to appear at separate locations that match the distance that the frame traveled (Özkan et al., 2021; Cavanagh 2020).

This perceived position shift occurs robustly across different visual environments such as apparent motion of a frame (instead of actual motion), whole background shifts instead of a moving frame (i.e., background pattern moving left to right), or 3D frame flipping motion (Cavanagh et al., 2022). In all cases, this frame-induced position shift yields a drastic perceptual effect, in some conditions resulting in 100% stabilization: The locations of dots were perceived in the frame’s coordinates as if the frame was not in motion at all. Such dramatic effects of perceived position shifts were taken as evidence that surrounding context (such as a moving frame) may act to stabilize the perception of relative locations during eye movements (Özkan et al., 2021).

But how and when does this perceived stability of object locations arise? Previous research has used electroencephalography (EEG) to examine the time course of when perceived (relative to physical) location processing arises in the visual processing stream in other paradigms that induce position shifts. For example, one study used the flash-grab effect, a motion-induced position shift where moving stimuli make flashing probes appear at projected locations and showed that illusory positions can be successfully decoded as early as 81 ms post-stimulus (Hogendoorn et al., 2015). Other investigations on the neural processes of this illusory position shift have used functional Magnetic Resonance Imaging (fMRI) and multivariate pattern analyses showing that activation patterns in early visual cortex carry information about the perceived locations (Kohler et al., 2017). Together, these data indicate that early visual processing is involved in coding the perceived position of objects. However, the recently discovered FIPS illusion is thought to be distinct from these motion-induced illusions, in particular because the magnitude of the perceived position shifts is not affected by the speed of the frame – which is different in other motion-induced illusions, such as the flash-grab effect. Instead, the critical factor that determines the FIPS illusion appears to be the distance that the frame travels from one side to the other (Özkan et al., 2021). Thus, it remains an open question of what the temporal dynamics are that underlie this frame-induced illusory position shift.

In the present study, to tease apart physical and perceived location processing during FIPS, we measured neural activity elicited by small black disks (‘probes’) that were presented either at the exact same physical location (along the central vertical midline) but *perceived* at different, peripheral locations due to a horizontally moving frame, and compared this to probe-elicited neural activity of stimuli that were physically at separate peripheral locations but matched the illusory perceived locations. Our results reveal that early visual-evoked potentials (VEPs) elicited by the probes were sensitive to the physical locations of stimuli starting at ∼70ms, as expected from previous research (e.g., Woldorff et al., 1997), while later VEPs starting at ∼140ms showed sensitivity to both perceived and physical location information. Using single-trial multivariate analyses, we further honed in on these effects to test whether similar neural activity patterns underlie these univariate effects. Indeed, we found that the probe locations could be reliably decoded across physical and perceived locations during the later time interval (140-180ms) but not during the earlier time interval (70-110ms). This suggests that the perception of spatial location arises around 140ms and that the visual processing of the illusory position shares neural activity patterns with the matched physical location during this later time interval.

## Results

### Behavior: Assessing individual illusion magnitudes before and after the EEG session

First, we assessed the magnitude of the perceived position shifts in all participants. Participants viewed flashing black disks that were surrounded by a frame that continuously moved back and forth from one side to the other (Figure 1, left bottom). After watching this stimulus, participants were asked to adjust the position of two disks so that they would match the distance between the previously seen disks. This method of adjustment allowed us to estimate the magnitude of the perceived position shift in each individual participant (for more details, see Supplemental Material). Illusion magnitudes were calculated by averaging the distance between the two response disks across three blocks of the behavioral session. Participants completed this adjustment task pre- and post the EEG session to ensure that participants would still perceive the illusion after being exposed to it for a longer time period. The average illusion magnitude for the pre-EEG sessions was 6.36° of visual angle. Interestingly, the average post-EEG session illusion magnitude was significantly reduced to 4.92° (t(16)=3.84, p=0.001; Cohen’s d_z_=0.61). Most importantly for the current study, both pre- and post-EEG session illusion magnitudes were reliably above 0° (pre-EEG: t(16)=13.89, p<0.001; post-EEG: t(16)=8.12, p<0.001) demonstrating that participants persistently saw the illusion before and after the EEG session despite the reduction in the magnitudes.

**Figure 1.**
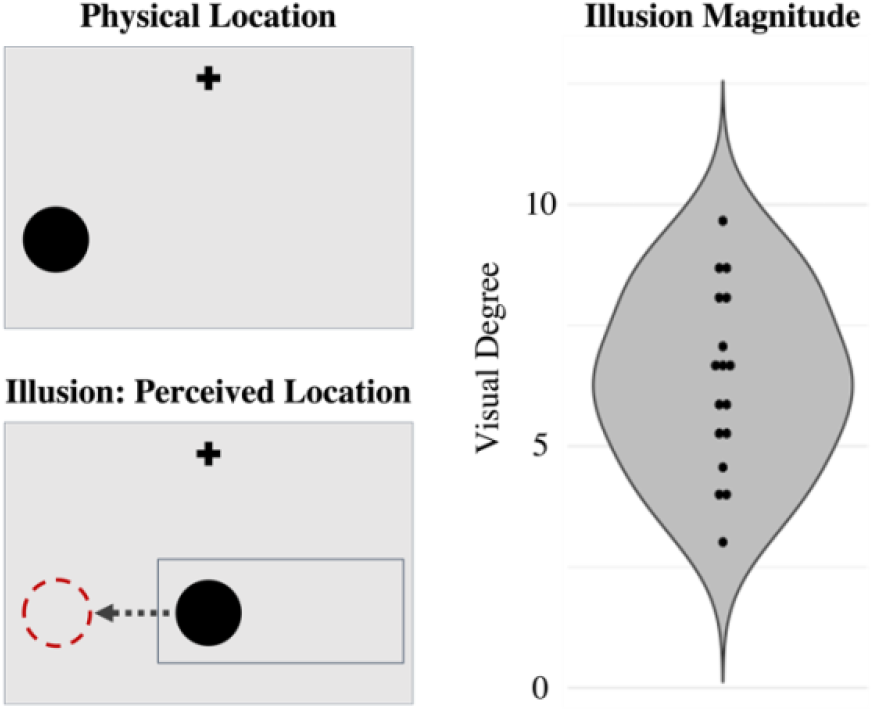
Illustration of the task displays (left) and reported magnitude of the illusory position shift (right). When a horizontally moving frame is shown (bottom left), a flashing dot that is presented centrally appears shifted towards the left or right sides. Using the reported illusion magnitude, we constructed the control condition where the dots are physically shifted the same amount (top left). Each participant’s pre-EEG session illusion magnitude is plotted on the right.

### ERPs: Early visually-evoked potentials for physical and perceived stimulus locations

There were three conditions in the EEG session: Perceived Location condition (disks are physically at the central location but perceived to appear on the left or right side due to a moving frame), Physical Location condition (disks are physically offset from the center matching the positions of each individual observer’s illusion magnitude), and Frame Only condition (only a moving frame is presented without the disks). For more details, see Supplemental Material.

Previous research has demonstrated that brain activity assessed via EEG differs across hemispheres for peripherally presented visual stimuli. Specifically, the P1 and N1 components of the event-related potential (ERP) have been found to be particularly sensitive to stimulus location, often occurring earlier in time and with larger amplitude when recorded over the occipital cortex contralateral to the stimulus location relative to ipsilateral (e.g., Luck & Hillyard, 1994; Woldorff et al., 1997). Thus, the main question of interest was whether we find similar lateralized effects for both the physical and perceived locations of the probe stimulus. Thus, we first examined the visually-evoked potential over occipital scalp sites that was elicited by the disks that were physically located on the left and right sides of the screen. Indeed, as can be seen in Fig. 2A, both the P1 (70ms-110ms) and N1 (140ms-180ms) components elicited larger amplitudes over the contralateral relative to ipsilateral occipital cortex with respect to the probe’s location.

**Figure 2.**
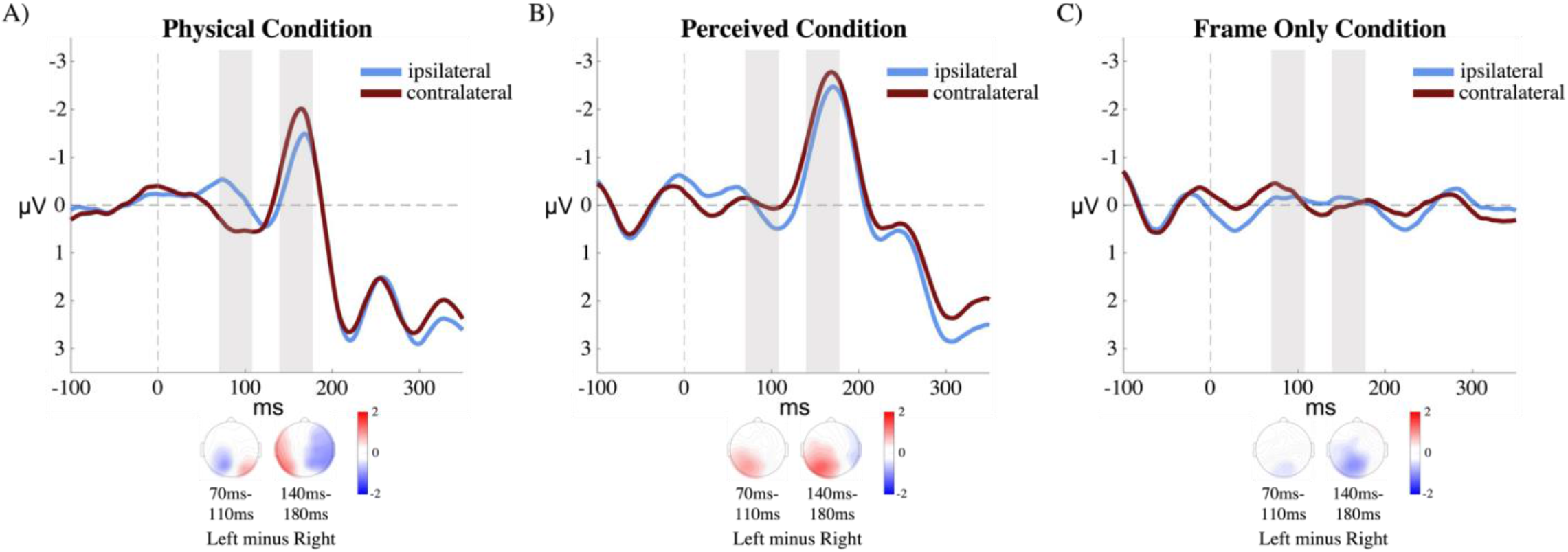
ERP results with the two time windows of interest highlighted in gray. **A)** In the physical condition, both P1 (70-110 ms) and N1 (140-180 ms) components show lateralized effects where amplitudes are larger over the hemisphere contralateral (red) relative to ipsilateral (blue) with respect to the disk location. **B)** In the perceived condition, only the N1 component shows a reliable lateralized effect, with the contralateral waveform showing a larger amplitude than ipsilateral. **C)** These effects cannot be explained by the presence of the moving frame as no lateralized activity was observed there.

Next, we examined whether the frame-induced perceived left and right locations generate a similar pattern in the ERP. Thus, we compared the waveforms over the hemisphere ipsilateral and contralateral with respect to the *perceived* probe location, although the physical locations were identical and non-lateralized along the vertical meridian (see Fig. 1 left bottom). Thus, if early P1 and N1 components are only sensitive to physical, but not perceived location information, we would expect to see no difference between these conditions. As can be seen in Fig. 2B, we observed no reliable amplitude difference between the waveforms recorded over the contralateral and ipsilateral hemisphere in the early P1 time period. Interestingly however, we observed a larger N1 amplitude over contralateral relative to ipsilateral occipital cortex, resembling the amplitude difference we observed in the physical stimulus condition (see Fig. 2).

These observations were confirmed statistically: A 2×2 within-subjects ANOVA with disk condition (physical vs. perceived) and laterality (contralateral vs. ipsilateral) as factors on the P1 amplitudes yielded a significant interaction (F(1, 17) = 35.476, p < 0.001). Pairwise comparisons using the t-tests with Bonferroni correction showed that the P1 amplitudes were significantly different in the physical condition but not in the perceived condition (physical condition P1: t(17) = 4.88, p < 0.001; perceived condition P1: t(17) = 1, p = 1.24). An 2×2 within-subjects ANOVA on the N1 amplitudes yielded a significant main effects of disk condition (F(1, 17) = 6.143, p = 0.024) and laterality (F(1, 17) = 10.719, p = 0.004), but no significant interaction (F(1, 17) = 0.842, p = 0.372, see Fig. 3).

**Figure 3.**
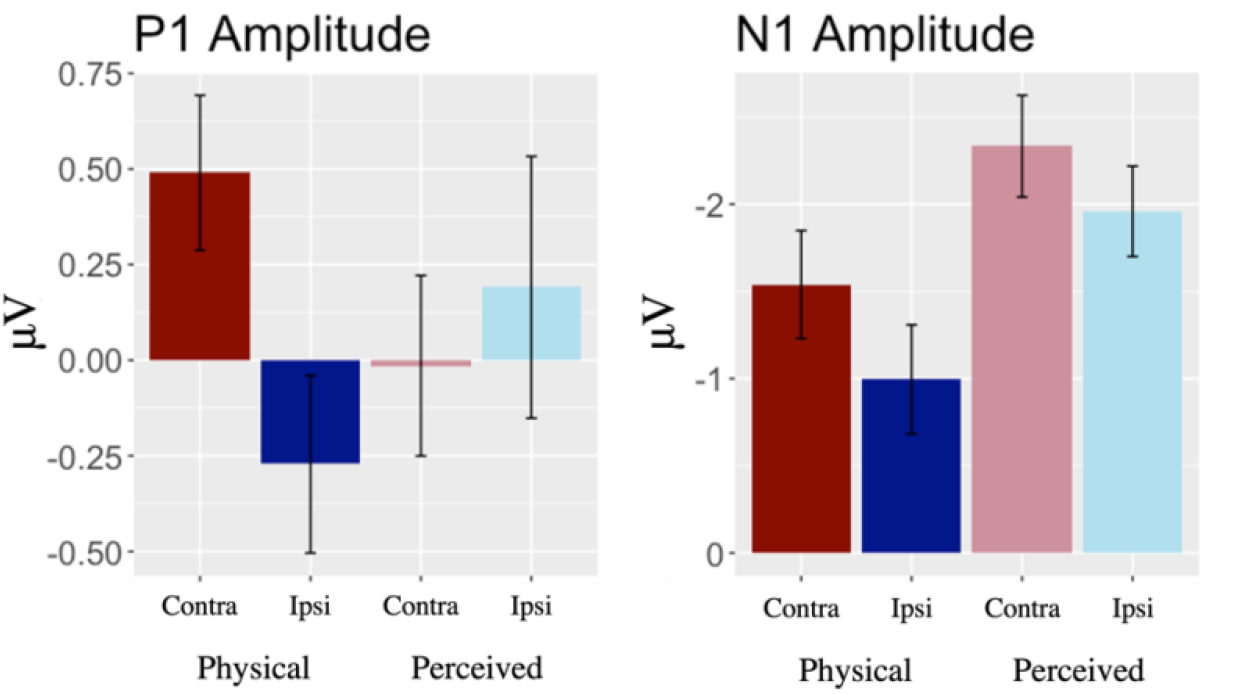
Mean amplitudes plotted for the P1 (left) and N1 (right) components. separately for the physical and perceived conditions. For the physical condition, both P1 and N1 show a reliable amplitude modulation such that activity recorded over contralateral occipital cortex is larger relative to activity recorded ipsilaterally. For the perceived location condition, only the N1 component shows this lateralized difference in amplitude.

**Figure 4.**
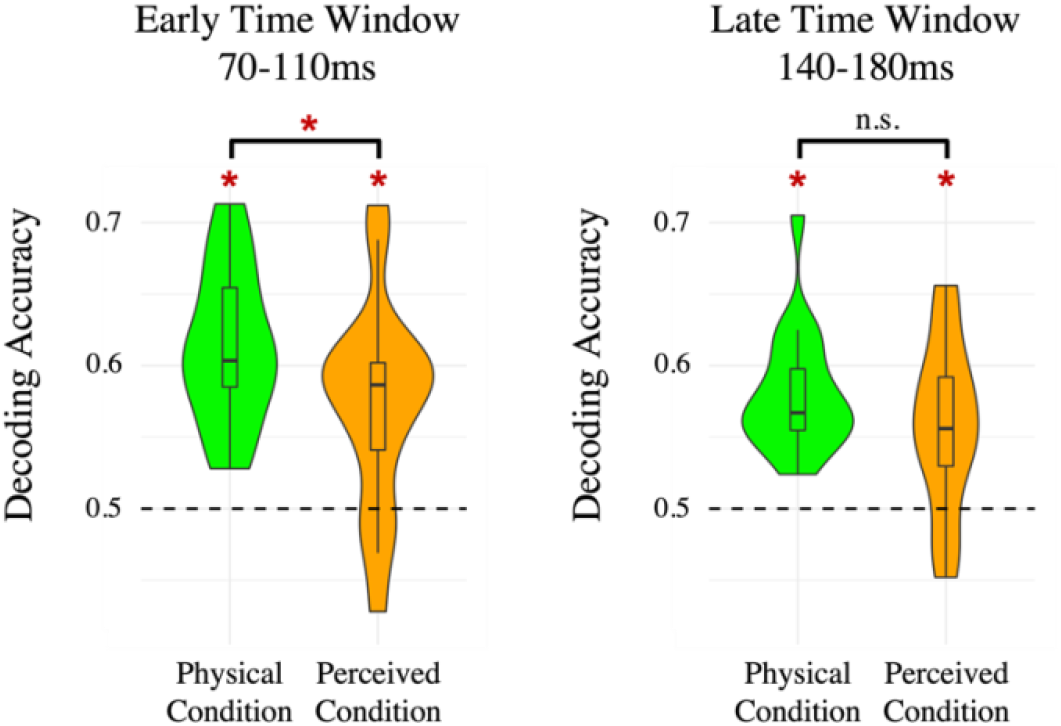
Within-condition decoding results for the early (left) and later (right) time windows. All conditions and time points resulted in above chance (50%) decoding performance. For the early time interval only, we observed higher decoding accuracy for the physical condition compared to perceived condition.

**Figure 5.**
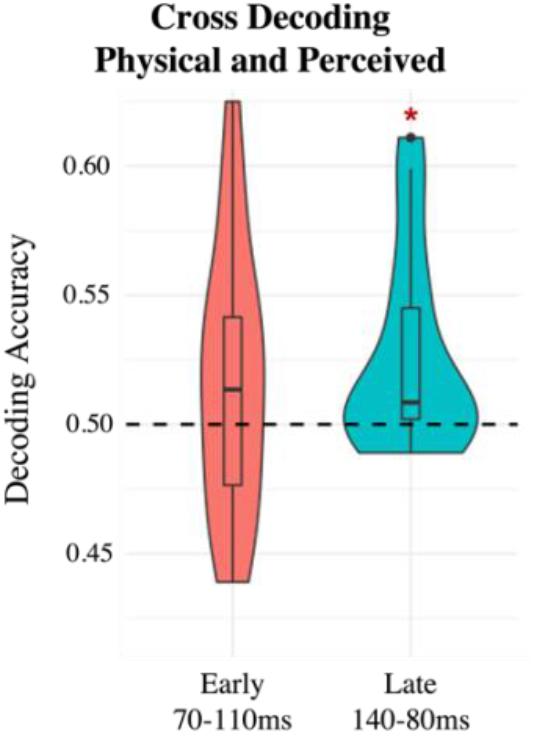
Cross-condition decoding results. We trained a classifier on the physical probe locations and tested whether it could reliably predict the perceived probe locations. Consistent with the late-arising position information for illusory locations observed in the ERP analysis, we find that we can only reliably cross-decode during the late time window.

In order to perceive the illusion, a continuously moving frame needs to be presented throughout the perceived location condition that was absent from the physical condition. To ensure that the differences in ERPs were not caused by the moving frame itself, we also analyzed the ERP amplitudes of the frame-only condition where there were no disks presented during the trial.

Because there were no disks to reference the laterality when computing the waveforms, we matched this analysis to the analysis used in the perceived condition where the frame being on the left side would imply a disk on the right side of the screen and vice versa. As expected, we found no clear visually-evoked potentials in the same P1 and N1 time windows and no differences between the contralateral and ipsilateral amplitudes in either time window (P1: t(17) = 0.89, p = 0.39; N1: t(17) = 0.77, p = 0.45; see Fig. 2C). Thus, the moving frame alone did not produce lateralized differences in the ERP.

### Neural Activity Patterns: Multivariate decoding analysis across physical and perceived stimuli

The ERPs provide clear evidence that perceived location information arises later in time than physical location information during FIPS. However, comparing the mean amplitudes of the ipsilateral vs. contralateral waveforms is a relatively coarse measure that tells us little about whether the neural processes underlying the coding of perceived and physical locations are different or shared. Thus, we extended our analyses to examine the neural activity patterns between processing the illusory positions versus the physical positions. To do so, we used a support vector machine (SVM) decoding approach using single-trial EEG activity in each condition. We focused this analysis on the two a priori defined time windows of interest that we also used in our ERP analyses. We reasoned that this would increase power and reduce the possibility of false positives, given that single trial data is relatively noisy. Furthermore, we were particularly interested to see whether the differences we observed in the univariate analyses would translate to multivariate decoding as this allows us to test whether the neural activity patterns for perceived and physical locations are shared or not. Thus, our decoding analysis focused on the a priori defined early (P1 component, 70ms-110ms) and late (N1 component, 140ms-180ms) time windows. We first established a baseline for how well we could decode the left versus right locations of the disks in the physical condition where disks were presented on the left and right side of the screen. We trained an SVM to distinguish between the left versus right disk locations based on single-trial EEG data averaged over each of these two time windows, and then tested this classifier on a leave-one-out hold-out set (Supplemental Material). On average across participants, we found decoding accuracy of 61.3% in the early time window and 58.1% in the later time window, both well above the chance level of 50% (early time window: t(17) = 8.99, p<0.001; late time window: t(17) = 8.06, p<0.001).

We then tested how well location information could be decoded in the perceived condition using the same approach. Interestingly, we found that decoding accuracies of disk location conditions (left vs. right) were significantly above chance level in both the early time window (mean performance 57.3%; t(17) = 4.37, p<0.001) and late time window (mean performance 58.1%; t(17) = 4.18, p<0.001). While the early time window resulted in above chance level decoding in both physical and perceived conditions, the physical location condition showed significantly higher decoding performance than the perceived location condition in this early time window (t(17) = 2.61, p=0.018). This was not the case for the late time window, in which case there was no significant difference in decoding performance between physical and perceived locations (t(17) = 1.65, p=0.12).

Using this decoding technique we can now ask: Are the illusory locations eliciting the same pattern of activity as the physical locations? We ran a cross-decoding analysis where we trained the model on the physical condition and tested on a random trial from the perceived condition. In the early time window this cross-decoding performance was not significantly above chance level (mean performance 51.6%; t(17) = 1.39, p=0.18). However, in the late time window cross-decoding accuracy was significantly above chance level (mean performance 52.5%, t(17) = 2.86, p=0.01), suggesting that the neural processes that give rise to physical and perceived locations are – at least in part – shared during the late time period.

## Discussion

Despite much research pointing to the importance of relational position coding to achieve visual stability, how the visual system accomplishes this – and at what stage of processing – remains not fully understood. Using a recently discovered visual illusion where a moving frame induces a paradoxical stabilization of probe positions (Özkan et al., 2021), we here show that relative position coding arises relatively late in the visual processing stream, but once reached, shares a similar neural code as position coding of objects that were not stabilized due to a frame.

We first examined visually-evoked potentials that were elicited by physical peripheral probes (with no frame present) or probes that matched these locations *perceptually* due to a moving frame, but were actually presented at the same physical position along the vertical midline. Our ERP findings showed a clear pattern: the earlier P1 component showed a lateralized effect only for the disks that were physically offset in location, with no reliable difference for perceptually shifted locations (frame-induced position shifts), while the later N1 component was sensitive to both physical and illusory disk locations. The P1 and N1 components are well established and long studied visually-evoked potentials known to reflect the early visual processes in occipital visual areas (e.g., Heinze et al., 1990; Hillyard & Anllo-Vento, 1998; Luck et al., 1994; Mangun, 1995, McDonald et al., 2013). The P1 component is typically interpreted as reflecting the initial feedforward sweep of visual processing, while the N1 component is thought to reflect both feedforward and feedback processing from higher visual and parietal areas. Consistent with this, prior studies have shown that dipoles that are underlying the P1 component are localized to various areas of the striate and extrastriate cortex (i.e., V3, V3a, middle occipital gyrus, and fusiform gyrus), while N1 generators are much more complex, distributed over more diverse areas including extrastriate cortex and also centro-parietal areas (e.g., Eimer, 1998; Hashimoto et al., 1999; Hoshiyama et al., 2001; Nakamura et al., 2000; Brecelj et al., 1998; Seki, 1996; Russo et al., 2002; Sun et al., 2022). Thus, our results suggest that physical location information is coded rapidly in the initial feedforward sweep of visual information processing (as indexed by the P1), but that later perceptual coding of information – here, the frame-induced stabilization of object locations, arises later in time (as reflected by the N1 component). This timing difference between physical versus illusory information may arise from more recurrent feedback and computations required to achieve visual stability via relational coding.

One potential concern in our design is that P1 and N1 amplitudes could be influenced by the physical difference between the displays of the two conditions: To induce the illusion, a continuously moving frame had to be presented which was not present in the physical condition. While neither the motion nor the location of the frame would necessarily predict larger P1 or N1 amplitudes at the contralateral side of the perceived disk location, we also included the moving frame-only condition where no disks were presented on the screen to ensure that the differences in ERP amplitudes were indeed not driven by the presence of the frame. This control condition did not result in any lateralized activity at the two time windows of interests; thus, it is unlikely that the frame itself affected the amplitude differences of the probe-evoked P1/N1 components in the illusory condition. To look into this further, we also examined the overall amplitudes of the P1 and N1 components across the two conditions and found no difference for the P1, but a reliable difference in amplitude for the N1 component, which was overall higher for the perceived condition compared to the physical condition. This may suggest that the processing of the illusory locations may require additional neural processes that are not present in the physical condition. However, there was no significant interaction of the lateralized effect across the two conditions, indicating that the main comparison of the contralateral versus ipsilateral N1 amplitudes was not affected by the overall difference across the conditions.

Using the multivariate decoding analysis, we further investigated whether the neural pattern of activity is sensitive to the physical and illusory locations across different time windows. Results showed that both physical and illusory locations could successfully be decoded during the early time window (70-110 ms) and the late time window (140-180 ms). Interestingly, we also found that decoding performance was significantly higher during the late time window relative to the early time window for the illusory condition only. One explanation of this result could be that the constant presentation of the moving frame during the illusory condition was – at least in part --contributing to the decoding results. To directly compare how the neural patterns compare between processing of the physical and illusory locations, we examined how well the model generalizes from the physical condition to the perceived condition. Results showed that cross-decoding between the two conditions was above chance level during the late time window, but not during the early time window. This suggests that the illusory positions share a neural code with the physical positions later on in time, and that early on in visual processing, these positions are dissociable by the patterns of electrical brain activity.

Our results relate to other studies that investigated the electrophysiological basis of illusory position shifts. In particular, Hogendoorn et al. (2015) examined electrophysiological activity during the motion-induced position shift, a form of position shift that is affected by motion of the stimuli, and found that cross-decoding between physical and illusory positions can be successful as early as 81 ms after the stimuli onset, a time point that’s analogous to the early time window in our analysis. The different results from Hogendoorn et al. (2015) and the current results where we find above chance level cross-decoding only during the late time window can be explained by the differences in the mechanisms underlying motion-induced position shift and FIPS. Behaviorally it has been shown that unlike the motion-induced position shift, the FIPS is not affected by low-level stimulus factors such as the speed of the motion or the location of where the motion energy is. It could be that the motion-induced position shift relies more on the low-level visual processes while the FIPS requires higher level processes that result in shifting the relevant electrophysiological response to a later time point that reflects these illusory positions. It is also true that in the current paradigm the two conditions differ in their stimuli properties (i.e., one has motion energy, one doesn’t), and this might be impacting the decoding performance across the two conditions. However, this seems unlikely to be the main driver of the observed effects as such stimulus-level differences should impact both the early and late time windows, and we selectively find above-chance cross-decoding performance for the late time window. Future investigations could aim to control for the differences between the two conditions.

Most broadly, our results suggest that physical location information is coded in the initial feedforward sweep of visual information processing and that later recurrent processes are involved in the relative coding of object locations to support visual stabilization.

## Limitations of the Study

The illusion magnitudes we found in the current study are on average smaller in comparison to the previous reports that showed 100% stabilization (illusion magnitude matching the frame’s path length; Özkan et al., 2021). Smaller illusion magnitude in the current study may be explained by the presence of a fixation cross that is necessary in the EEG task to avoid eye movements. While previous investigation on the FIPS claim that having a fixation point does not eliminate the illusion (Özkan et al., 2021), having a concrete reference point that is also presented at the vertical meridian may produce reduction in the illusion magnitude. Importantly however, participants reported to consistently perceive the central disks shifted in location before and after the EEG session in the current study.

## Methods and Analysis (Supplemental Material)

### Participants

Eighteen participants were recruited from Dartmouth College, all reported normal or corrected-to-normal vision and were between 18 and 28 years of age. All participants gave informed consent as approved by the Committee for the Protection of Human Subjects at Dartmouth College and were paid $20/hr or 1 class credit/hr for their time. All participants included in the study reported seeing the illusion on the demo video, however, three participants who did not show more than 2° of visual angle of illusion on the initial behavioral session were excluded from the study. The value of one participant’s illusion magnitude was not saved correctly, resulting in 17 total participants for the behavioral analysis.

### Stimuli and Apparatus

Participants were seated approximately 37 cm in front of a 24-in computer monitor (1920×1080) in a dark, electrically shielded chamber. Stimuli were presented on a white background via the Psychophysics Toolbox (Brainard, 1997; Pelli 1997) in MATLAB. In both the behavioral and EEG sessions, a small black fixation cross (0.4° x 0.4° of visual angle) was presented at 7° above from the center of the screen. On each trial, a rectangular gray frame (18.5° width x 3° height x 0.15° thickness; exact color value varied across participants, average [RGB: 99 99 99]) was presented at 12° below the center of the screen. During the trial, this frame moved left and right at a speed of 2.6° (60 pixels) per frame within the bounded range of 30° left and right from the center. When the frame made contact with the left or right border, a faint gray or black disk (2.4° radius) appeared 12° below the center of the screen. For the EEG session, these disks were presented on the vertical meridian (Perceived Location condition), or their positions were individually adjusted towards the left/right sides of the screen based on the individual’s magnitude of the illusion (Physical Location condition). In the behavioral part of the experiment, participants only completed the illusory trials (Perceived Location condition) and, after each trial, were instructed to adjust the distance between two response disks to match the locations that they had perceived during that trial (see *Experimental Procedure*).

### Experimental Procedure

Participants first completed the behavioral session and then went on to the EEG session. In the behavioral session, the magnitude of the FIPS illusion was measured for each participant. Participants were instructed to maintain their gaze at the fixation cross located at the top of the screen throughout the entire duration of each trial. On each trial, a frame was presented at the bottom of the screen and continuously moved horizontally left and right along a preset path at the speed of 2.6° per 8.3 ms. The frame stopped when it reached the left or right bound of the path, and a disk appeared at the bottom of the screen on the vertical meridian for 100 ms. Each trial consisted of 10 disks flashing. In most cases the disks were a faint gray color ([RGB: 190 190 190]) that mostly blended in with the background. At two random intervals, a black disk was presented: one when the frame reached the left edge, and one when the frame reached the right edge. We used these different disk colors to avoid overlap in the ERPs elicited by the disks during the EEG session while maximizing the illusion magnitude (see more details below). After the illusion display, participants reported the locations of the previously seen two black disks by adjusting the locations of two response probes. These response probes were first presented at the vertical meridian (at the bottom of the screen at center width), just as during the illusion itself, and as participants pressed the ‘c’ key to move them further apart or the ‘m’ key to move them closer together. When participants felt confident that the response probes were adjusted to reflect the perceived distance between the two black disks, participants pressed the spacebar and moved onto the next trial. Participants completed 12 trials per block and repeated 3 blocks. Illusion magnitudes were averaged across all three blocks.

The EEG session consisted of 3 conditions: Perceived Location condition (disk+frame), Physical Location condition (disk-only), and Frame Only condition. The display in the EEG session was identical as in the behavioral session: In the Perceived Location condition a moving frame and disks were presented every time the frame hit either left or right edge of the preset path. Dots were presented 10 times per trial with most of them being faint gray placeholders and only 2 or 3 black dots at random intervals. The black disks were the critical stimuli for the EEG analysis: ERPs were time-locked to these high-contrast black disks. The faint gray disks were presented with the sole purpose to strengthen the perception of the illusion which is more apparent with a continuous stream of disk presentations, while not eliciting strong visually-evoked potentials due to their low contrast. In the Physical Location condition, only the black disks were presented at the locations of each individual’s illusion magnitude. This means that if the participant reported 5° of illusion, then the disks were presented 5° left or right from the center width. This effectively simulates the illusory locations of each of our participants. The faint gray disks were omitted in the disk-only condition in order to eliminate the expectancy of the disk location based on oscillatory disk presentation. In the Frame Only condition, only the moving frame and faint gray placeholder disks were presented. At the intervals where the black disks would have shown up, no disks were presented on the screen. After each trial participants reported how many black disks were presented during the trial (0, 2, or 3) and received feedback on their response. The purpose of the task was to ensure that participants would be attending to the disks at the bottom of the screen. Participants completed the task until the end of the scheduled session, which resulted on average of 602 total trials per participant (ranging from 488 to 710 trials). Average accuracy of the disk-counting task was 98% correct.

After the EEG session, three more blocks of the behavioral session were run to compare how the illusion magnitudes changed over time.

### Electrophysiological Recordings and Analysis

EEG was recorded continuously from 32 Ag/AgCl electrodes mounted in an elastic cap and amplified by an ActiCHamp amplifier (BrainProducts, GmbH). Electrodes were arranged according to the 10–20 system. The horizontal electrooculogram (HEOG) was recorded from two additional electrodes placed on the external ocular canthi which were grounded with an electrode placed on the neck of the participant. All scalp electrodes were referenced to the right mastoid online and were digitized at 500 Hz. EEG data was filtered with a bandpass of 0.01-112.5 Hz online.

Data processing was performed using EEGLAB (Delorme & Makeig, 2004) and ERPLAB (Lopez-Calderon & Luck, 2014) toolboxes in MATLAB (The MathWorks, Natick, MA) and custom-written scripts. Data were epoched from -100 ms to 450 ms with respect to the onset of the black disk (or the equivalent time period for the frame only condition). A semiautomatic procedure was performed to remove any EEG epochs that were contaminated by artifacts such as eye movements and blinks. For each participant, each epoch was visually inspected to check the automated procedure. Artifact-free data were re-referenced to the average of the left and right mastoids.

#### ERP analysis

For the ERP analysis, waveforms were averaged separately with respect to the left and right disk locations and were then collapsed across disk location and hemisphere of recording to obtain waveforms recorded ipsilaterally and contralaterally relative to the physical or perceived locations. For the frame-only condition, the same procedure was performed as for the perceived location condition where the dot location was assumed to be left or right with respect to the frame location. ERPs were low-pass filtered (half-amplitude cutoff at 25Hz; slope of 12dB/octave) to remove high-frequency noise. Mean amplitudes for each participant and condition were measured with respect to a 100-ms prestimulus period (−100 to 0 ms from disk onset, or equivalent time period for frame only condition). The exact time windows and electrode sites for each ERP analysis were chosen a priori based on previous research and matched across all analyses. Our main analysis was focused on the neural activity during the P1 (70ms-110ms) and N1 (140ms-180 ms) time windows. Both ERP components were measured at the same eight parietal-occipital electrode sites (O1/O2/PO3/PO4/PO7/PO8/P7/P8). Amplitudes of ipsilateral and contralateral ERPs were then compared using 2×2 ANOVA with conditions and laterality as factors, followed up by pairwise t-tests for significant interactions.

#### Multivariate decoding analysis

For the decoding analysis, we used single-trial EEG activity averaged either over the early time window of the P1 (70ms-110ms) and the later time window of the N1 component (140 ms-180 ms). We trained a classifier to distinguish between the left and right locations of the stimulus based on a support vector machine (SVM) decoding approach (Cox & Savoy, 2003). The input feature vector in each case consists of 31 features: averaged activity of the 40-ms time window from each of 31 electrodes. We then applied a leave-one-out classification procedure with 500 iterations where a randomly chosen trial from each condition (left or right) is left out. We applied this method separately for each time window and condition: Perceived Location (disk+frame), Physical Location (disk-only), and Frame Only. Then, to test whether the neural activity pattern is similar across the conditions, we tested whether training in one condition would generalize to another condition. Thus, we trained a model on the disk-only condition (physical location condition) and tested on a random trial of the disk+frame condition (perceived location condition). Performance was then averaged over the 500 iterations for each participant. In each condition, we then performed one-sample t-tests versus chance performance (50%) to assess the statistical significance of the decoding accuracy.

## References

Asch, S. E., & Witkin, H. A. (1948). Studies in space orientation; perception of the upright with displaced visual fields. Journal of Experimental Psychology. 38, 325–337.

Brainard, D. H., (1997). The psychophysics toolbox. Spatial vision, 10(4), 433–436.

Brecelj, J., Kakigi, R., Koyama, S., & Hoshiyama, M. (1998). Visual evoked magnetic responses to central and peripheral stimulation: Simultaneous VEP recordings. Brain Topography, 10(3), 227–237.

Cavanagh, P., Anstis, S., Lisi, M., Wexler, M., ‘t Hart, M., Shams-ahmar, M., & Saleki, S. (2022). Exploring the Frame Effect. Journal of Vision 22(12):5, 1-13.

Cavanagh, P. (2020). Frame-induced position shifts. Journal of Vision, 20, 607.

Cox D. D., & Savoy R. L. (2003). Functional magnetic resonance imaging (fMRI) “brain reading”: detecting and classifying distributed patterns of fMRI activity in human visual cortex. Neuroimage 19:261–270.

Delorme, A., & Makeig, S. (2004). EEGLAB: an open source toolbox for analysis of single-trial EEG dynamics including independent component analysis. Journal of Neuroscience Methods, 134(1): 9–21.

Eimer, M. (1998). Mechanism of Visuospatial Attention: Evidence from Event-related Brain Potentials. Visual Cognition, 5(1/2), 257–286.

Hashimoto, T., Kashii, S., Kikushi, M., Honda, Y., Nagamine, T., & Shibasaki, H. (1999). Temporal profile of visual evoked responses to pattern-reversal stimulation analyzed with whole-head magnetometer. Experimental Brain Research, 125, 375–382.

Heinze, H. J., Mangun, G. R., & Hillyard, S. A. (1990). Visual event-related potentials index perceptual accuracy during spatial attention to bilateral stimuli. Psychophysiological brain research, 196–202.

Hillyard, S. A., & Anllo-Vento, L. (1998). Event-related brain potentials in the study of visual selective attention. Proceedings of the National Academy of Sciences, 95, 781–787

Hogendoorn, H., Verstraten, F. A. J., & Cavanagh, P. (2015). Strikingly rapid neural basis of motion-induced position shifts revealed by high temporal-resolution EEG pattern classification. Vision Research, 113(PA), 1–10. https://doi.org/10.1016/j.visres.2015.05.005

Hoshiyama, M., & Kakigi, R. (2001). Effects of attention on pattern-reversal visual evoked potentials: Foveal field stimulation versus peripheral field stimulation. Brain Topography, 13(4), 293–298. https://doi.org/10.1023/A:1011132830123

Kohler, P. J., Cavanagh, P., & Tse, P. U. (2017). Motion-induced position shifts activate early visual cortex. Frontiers in Neuroscience 11: 168.

Lopez-Calderon, J., & Luck, S. J. (2014). ERPLAB: An open-source toolbox for the analysis of event-related potentials. Frontiers in Human Neuroscience, 8, Article 213. https://doi.org/10.3389/fnhum.2014.00213

Luck, S. J., & Hillyard S. A. (1994) Spatial filtering during visual search: evidence from human electrophysiology. Journal of Experimental Psychology: Human Perception and Performance 20:1000–1014.

Luck, S. J., Hillyard, S. A., Mouloua, M., Woldorff, M. G., Clark, V. P., & Hawkins, H. L. (1994). Effects of Spatial Cuing on Luminance Detectability: Psychophysical and Electrophysiological Evidence for Early Selection. Journal of Experimental Psychology: Human Perception and Performance 20(4): 887–904.

Mangun G. R., & Hillyard, S. A. (1995). Mechanisms and models of selective attention. Electrophysiology of mind: Event-related potentials and cognition, 40–85.

McDonald, J. J., Störmer, V. S., Martinez, A., Feng, W., & Hillyard, S. A. (2013). Salient sounds activate human visual cortex automatically. Journal of Neuroscience, 33(21), 9194–9201.

Morgan, M., Grant, S., Melmoth, D., & Solomon, J. A. (2015). Tilted frames of reference have similar effects on the perception of gravitational vertical and the planning of vertical saccadic eye movements. Experimental Brain Research. 233, 2115–2125.

Nakamura, M., Kakigi, R., Okusa, T., Hoshiyama, M., & Watanabe, K. (2000). Effects of check size on pattern reversal visual evoked magnetic field and potential. Brain Research, 872(1– 2), 77–86. https://doi.org/10.1016/S0006-8993(00)02455-0

Özkan, M., Anstis, S., ‘t Hart, B. M., Wexler, M., & Cavanagh, P. (2021). Paradoxical stabilization of relative position in moving frames. Proceedings of the National Academy of Sciences, 118(25), 1–8.

Pelli, D. G. (1997). The VideoToolbox software for visual psychophysics: transforming numbers into movies. Spatial vision

Post, R. B., Welch, R. B., & Whitney, D. (2008). Egocentric and allocentric localization during induced motion. Experimental Brain Research. 191, 495–504.

Russo, F. D., Martínez, A., Sereno, M. I., Pitzalis, S., & Hillyard, S. A. (2002). Cortical sources of the early components of the visual evoked potential. Human Brain Mapping, 15(2): 95–111.

Seki, K. (1996). Neuromagnetic evidence that the P100 component of the pattern reversal visual evoked response originates in the bottom of the calcarine fissure. Electroencephalography and Clinical Neurophysiology/Evoked Potentials Section, 100(5), 436–442. https://doi.org/10.1016/s0921-884x(96)95098-5

Sun, R., Sohrabpour, A., Worrell, G., & He, B. (2022). Deep neural networks constrained by neural mass models improve electrophysiological source imaging of spatiotemporal brain dynamics. Proceedings of the National Academy of Sciences of the United States of America, 119(31).

Wertheim, A. H. (1981). On the relativity of perceived motion. Acta Psychologica, 48(1-3), 97– 110.

Woldorff, M. G., Fox, P. T., Matzke, M., Lancaster, J. L., Veeraswamy, S., Zamarripa, F., Seabolt, M., Glass, T., Gao, J. H., Martin C. C., & Jerabek P. (1997). Retinotopic organization of early visual spatial attention effects as revealed by PET and ERPs. Human Brain Mapping, 5(4):280–6.

